# Hetero-oligomeric CPN60 resembles highly symmetric group I chaperonin structure revealed by Cryo-EM

**DOI:** 10.1101/432013

**Authors:** Qian Zhao, Xiang Zhang, Frederik Sommer, Na Ta, Ning Wang, Michael Schroda, Yao Cong, Cuimin Liu

**Affiliations:** State Key Laboratory of Plant Cell and Chromosome Engineering, Institute of Genetics and Developmental Biology, Chinese Academy of Sciences, University of Chinese Academy of Sciences, Beijing, 100101, China.; National Center for Protein Science Shanghai, State Key Laboratory of Molecular Biology, CAS Center for Excellence in Molecular Cell Science, Shanghai Institute of Biochemistry and Cell Biology, Chinese Academy of Sciences, University of Chinese Academy of Sciences, Shanghai, China 201210; Shanghai Science Research Center, Chinese Academy of Sciences, Shanghai, China 201210; Molecular Biotechnology and Systems Biology, TU Kaiserslautern, Erwin-Schroedinger Str. 70, 67663 Kaiserslautern, Germany

**Keywords:** Chaperonin, Rubisco, Protein folding, Photosynthesis, CPN60

## Abstract

The chloroplast chaperonin system is indispensable for the biogenesis of Rubisco, the key enzyme in photosynthesis. Using *Chlamydomonas reinhardtii* as the model system, we revealed that chloroplast chaperonin is consisted of CPN60α, CPN60β1, and CPN60β2, and co-chaperonin is composed of three subunits CPN20, CPN11 and CPN23 *in vivo*. CPN20 homo-oligomers and all possible other chloroplast co-chaperonin hetero-oligomers are functional, but only CPN11/20/23-CPN60αβ1β2 pair can fully replace GroES/GroEL in *E. coli* at stringent growth condition. Endogenous CPN60 was purified and its stoichiometry was determined to be 6:2:6 for CPN60α:CPN60β1:CPN60β2. The cryo-EM structures of endogenous CPN60αβ1β2/ADP and CPN60αβ1β2/co-chaperonin/ADP were solved at resolutions of 4.06 Å and 3.82Å, respectively. In both hetero-oligomeric complexes the chaperonin subunits within each ring are highly symmetric. The chloroplast co-chaperonin CPN11/20/23 formed seven GroES-like domains through hetero-oligomerization which symmetrically interact with CPN60αβ1β2. Our structures also reveal an uneven distribution of roof-like structures in the dome-shaped CPN11/20/23 and potentially diversified surface properties in the folding cavity of CPN60αβ1β2 that might enable the chloroplast chaperonin system to assist in the folding of specific substrates.

## Introduction

Cellular protein homeostasis is maintained by a complex network of molecular chaperones (Bukau et al., 2006; Hartl et al., 2011; Lee and Tsai, 2005; Ramos, 2011; Saibil, 2013). Group I chaperonins are widely present in bacteria (GroEL) and endosymbiotic organelles of eukaryotes such as chloroplasts (Cpn60) and mitochondria (Hsp60) (Horwich, 2013; Yebenes et al., 2011). Biochemical and structural studies on the prototypical GroES/GroEL system from *Escherichia coli* has shed substantial light on the molecular mechanisms underlying group I chaperonin-assisted protein folding (Hayer-Hartl et al., 2016). GroEL is a large cylindrical complex, composed of two homo-oligomeric heptameric rings which are stacked together back to back (Braig et al., 1994). GroEL functionally cooperates with co-chaperonin GroES in an ATP-dependent manner to provide a central cavity where protein substrates are isolated from the crowded cellular environment and can fold correctly (Harris et al., 1994; Hayer-Hartl et al., 2016; Xu et al., 1997). GroEL subunits consist of three structurally and functionally distinct domains: the equatorial domain binds ATP and mediates almost all inter-ring and intra-ring communications. The apical domain binds nonnative substrate proteins and GroES. The hinge-like intermediate domain connects the other two domains and transmits the allosteric signal triggered by ATP binding and hydrolysis (Xu et al., 1997). The co-chaperonin GroES assembles into a dome-shaped heptameric ring and interacts with GroEL through mobile loops (Hunt et al., 1996; Xu et al., 1997). The interaction between GroES and GroEL heptamers drives the conformational change of GroEL that results in a protein folding nano-cage with two-fold increased volume (Chaudhry et al., 2003; Xu et al., 1997). Moreover, the physical environment of the nano-cage formed by chaperonin and co-chaperonin is crucial for the folding pathway of certain protein substrates (Clare et al., 2006; Tang et al., 2006).

Unlike the homo-oligomeric chaperonin systems from bacteria (GroES/GroEL) and mitochondria (Hsp10/Hsp60), the chloroplast chaperonin system is much more complicated, as all genomes of photosynthetic eukaryotes encode multiple copies of chloroplast chaperonin and co-chaperonin genes (Hill and Hemmingsen, 2001; Trosch et al., 2015; Vitlin Gruber et al., 2013; Weiss et al., 2009; Zhao and Liu, 2017). Chloroplast chaperonins harbor two subunit subtypes, termed Cpn60α and Cpn60β, which share about 50% sequence identity. Chloroplast co-chaperonin subunits are also divided into two subtypes, the conventional ~10 kDa GroES-like Cpn10 and the ~20 kDa Cpn20 that consists of two tandem Cpn10 domains joined head-to-tail (Baneyx et al., 1995; Bertsch et al., 1992; Musgrove et al., 1987). Both chloroplast chaperonin and co-chaperonin complexes are liable to exist as intricate hetero-oligomers even though Cpn60β and Cpn20 homo-oligomers have been shown to be functional *in vitro* (Bai et al., 2015; Dickson et al., 2000; Tsai et al., 2012; Viitanen et al., 1995; Vitlin Gruber et al., 2014). Another unique aspect of the chloroplast chaperonin system compared to its bacterial and mitochondrial counterparts is the capability to assist in the folding and assembly of ribulose 1,5-bisphosphate carboxylase/oxygenase (Rubisco), which accounts for ~30%-50% of soluble protein in chloroplasts (Barraclough and Ellis, 1980; Dhingra et al., 2004). A recent study has shown that the Cpn60αβ hetero-oligomer is indispensable for the functional assembly of *Arabidopsis thaliana* Rubisco in *Escherichia coli*, while the chloroplast co-chaperonin Cpn20 can be substituted by GroES (Aigner et al., 2017). It was proposed that the diversified chloroplast chaperonin system has adapted to accommodate specific substrates. Genetic studies have shown that the C-terminal amino acids of the Cpn60β4 subunit in the Cpn60αβ hetero-oligomer from *Arabidopsis* are specifically required for the folding of chloroplast proteins NdhH. Moreover, KASI, a protein important for the formation of the heart-shaped *Arabidopsis* embryo, depends on Cpn60α2 for proper folding (Ke et al., 2017; Peng et al., 2011). However, the biochemical and structural features responsible for these specialized functions of the hetero-oligomeric chloroplast chaperonin system remain elusive.

*Chlamydomonas reinhardtii* was regarded as well-suited for investigating the mechanism of chloroplast chaperonins, as its genome encodes fewer chaperonin and co-chaperonin isoforms when compared to land plants (Schroda, 2004). *Chlamydomonas* encodes three chloroplast chaperonin subunits, CPN60α, CPN60β1 and CPN60β2，as well as three co-chaperonin subunits that according to their molecular masses were named CPN11, CPN20 and CPN23 (Schroda, 2004). An analysis of the ability of individual *Chlamydomonas* CPN60 subunits to assemble into oligomers revealed that CPN60α was incapable of self-oligomerizing, but could form functional hetero-oligomers with CPN60β (Bai et al., 2015). Furthermore, domain swappings between CPN60α and CPN60β demonstrated that the CPN60α apical domain could not functionally cooperate with co-chaperonin GroES, but recognized its cognate substrate, the Rubisco large subunit from *Chlamydomonas,* more efficiently than the CPN60β apical domain and vice versa, suggesting one way of functional differentiation (Zhang et al., 2016b). Regarding chloroplast co-chaperonins it is not clear how the six/eight-fold symmetry realized in functional Cpn20 homo-oligomers matches with the heptameric chaperonin cylinder, although a recent biochemical study has claimed that a prefect match with the chaperonin was no absolute prerequisite for a functional interaction (Baneyx et al., 1995; Guo et al., 2015; Koumoto et al., 1999). Several *in vitro* studies have suggested that, similar to Cpn60s, the co-chaperonins might also form Cpn20-Cpn10 hetero-oligomers *in vivo* (Tsai et al., 2012; Vitlin Gruber et al., 2014). It appears that hetero-oligomerization of different subunits might not only be a smart way to solve the symmetry mismatch problem, but also might increase the flexibility of the chloroplast chaperonin system to provide adapted folding environments for specific substrates.

In this study we aimed at investigating the structure and stoichiometries of the native chaperonin-co-chaperonin complexes from *Chlamydomonas*. Biochemical assays showed that hetero-oligomeric co-chaperonin complexes containing CPN11, CPN20, and CPN23 were fully functional in cooperation with CPN60αβ1β2. Chaperonin CPN60αβ1β2 was purified from *Chlamydomonas* and the stoichiometry of the subunits was determined by mass spectrometry. By cryo-EM single particle analysis we determined the structure of CPN60αβ1β2 and of CPN60αβ1β2 incubated with recombinant CPN11/20/23. The overall structure and cavity size of the hetero-oligomeric chloroplast chaperonin system resemble that of its homo-oligomeric counterpart GroES/GroEL, but with potentially an asymmetric surface electrostatic potential.

## Results

### Chloroplast chaperonin CPN60 cooperates with its cognate co-chaperonin and complements GroEL/ES function

Previous studies have shown that hetero-oligomers consisting of Cpn10 and Cpn20 occurred frequently *in vitro*, which raised the question whether such hetero-oligomers represented functional cochaperonins also *in vivo* (Tsai et al., 2012; Vitlin Gruber et al., 2014). To confirm the existence of chloroplast co-chaperonin hetero-oligomers *in vivo*, we carried out co-immunoprecipitation assays from *Chlamydomonas* chloroplast stroma and total protein using an antiserum against CPN20. Precipitated proteins were separated by SDS-PAGE and the corresponding bands were analyzed by mass spectrometry (Figs. 1A and S1A). In addition to the three CPN60 subunits, CPN23 and CPN11 were also co-precipitated with CPN20, suggesting that chloroplast co-chaperonin subunits assemble into hetero-oligomers *in vivo*. Next we tested whether CPN11/20/23 heterooligomers are formed in *E. coli* and can interact with native chaperonins. For this, we expressed CPN11 with a hexa-histidine tag at its C-terminus together with CPN20 and CPN23 in *E. coli*. As shown in Fig. S1B, CPN20 and CPN23 could be co-purified with CPN11 on a nickel column, indicating that CPN11/20/23 heterooligomers are indeed formed in *E. coli*. Using the purified CPN11_6xHis_/20/23 complex as bait, CPN60α was co-purified from *Chlamydomonas* chloroplast stroma extracts only in the presence of ATP, indicating that CPN11/20/23 functionally interacts with native chloroplast chaperonin complexes (Fig. 1B).

**Figure 1.**
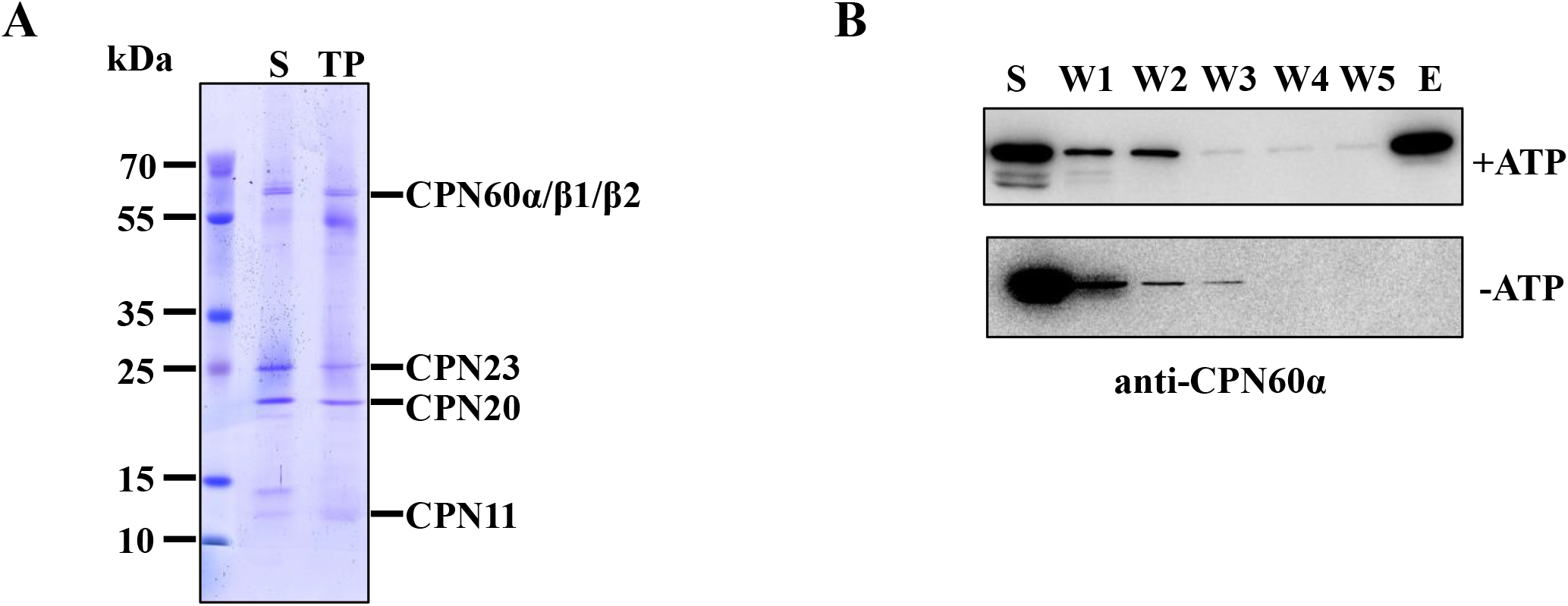
Composition of the chaperonin system in the chloroplast of *Chlamydomonas reinhardtii*.. **(A)** Stroma proteins (S) and soluble total proteins (TP) from 2 L *Chlamydomonas* cells were incubated with protein A–Sepharose coupled to anti-CPN20 serum. Co-immunoprecipitated proteins were separated on a 5–13% SDS-polyacrylamide gel and visualized by Coomassie blue staining. The identity of the marked proteins was verified by mass spectrometry. **(B)** Pull-down of endogenous CPN60α/β1/β2 by CPN11_6xHis_/20/23. 1 mg of recombinantly purified CPN11_6xHis_/20/23 was immobilized on Ni-NTA beads and incubated with stromal protein (S) from isolated chloroplasts from 1 L *Chlamydomonas* cells supplemented with or without 1 mM ATP for 4 h at 4°C, respectively. Proteins were eluted (E) with 0.5 M imidazole after five washing steps (W1-5) and were analyzed by immunoblotting using an antiserum against CPN60α.

To investigate the optimal co-chaperonin for chaperonin, more different combination of chaperonin/cochaperonin subunits was induced in *E.coli* MGM100, a conditionally GroES/GroEL-deficient strain, under normal (37°C) and stringent heat shock (45°C) growth conditions. When expressed in *E.coli* MGM100, CPN11 and CPN23 alone were unable to replace GroES when co-expressed with GroEL or CPN60αβ1β2, while CPN20 alone or any possible co-chaperonin combination (CPN11/20, CPN11/23 CPN20/23, and CPN11/20/23) could fully replace GroES at 37°C (Figs. 2A and S2). This indicates that a prerequisite for the functionality of CPN11 and CPN23 is their incorporation into co-chaperonin hetero-oligomers. However, GroEL with any of these co-chaperonins failed to support MGM100 growth at 45_o_C (Fig. 2A; Fig. S2A), while CPN60αβ1β2 with CPN20, CPN11/20, and CPN11/20/23 supported growth at 45°C, suggesting a species-specific functional cooperation between co-chaperonin and chaperonin (Figs. 2A and S2B).

**Figure 2.**
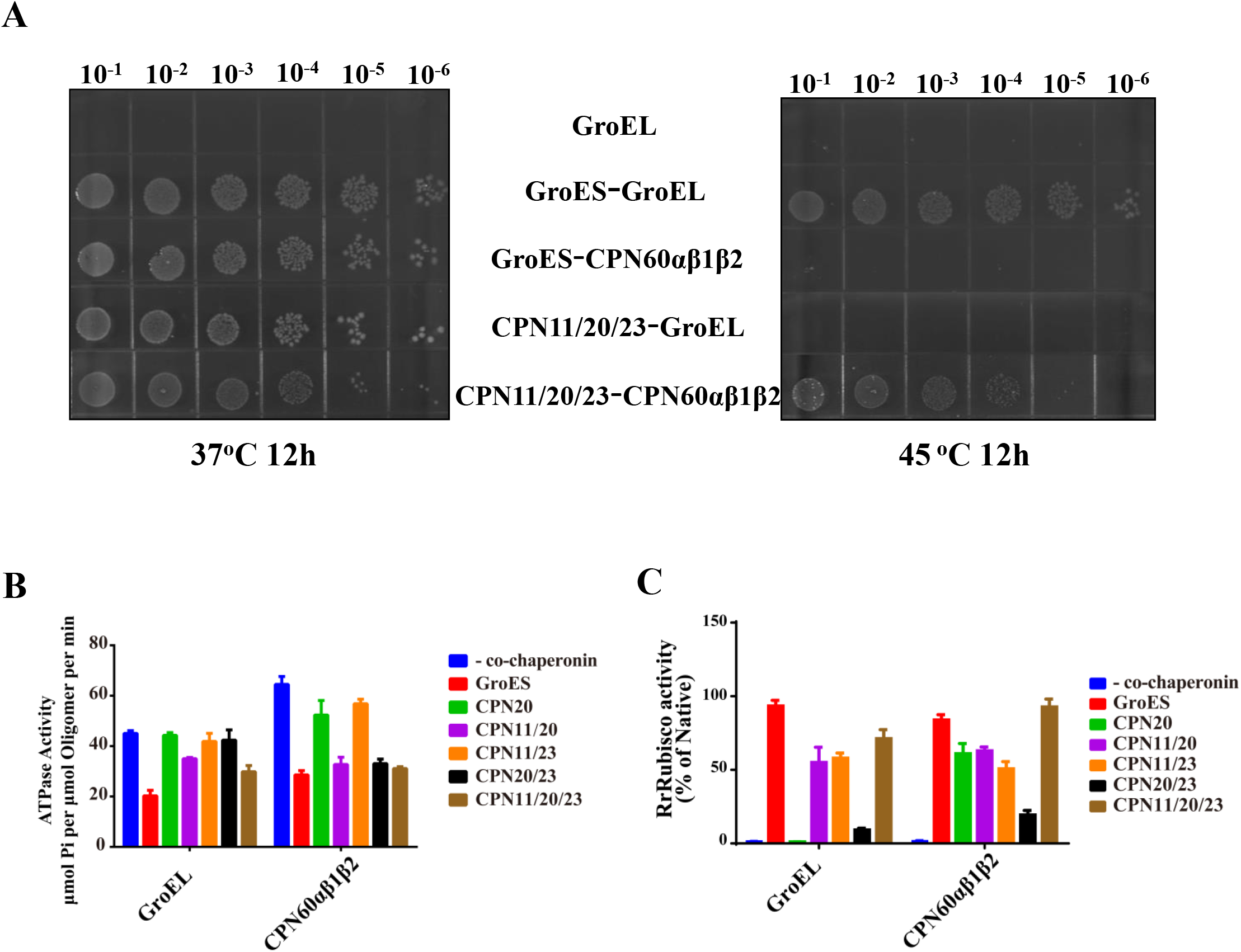
CPN11/20/23 form a functional chloroplast cochaperonin. **(A)** Complementation assays were carried out in *E. coli* strain MGM100, in which the native promoter of the endogenous GroE operon is replaced by the pBAD promoter. MGM100 was transformed with pQLinKT plasmids containing GroEL (negative control), GroES/GroEL (positive control), GroES-CPN60αβ1β2, CPN11/20/23-GroEL and CPN11/20/23-CPN60αβ1β2. Ten-fold-serial dilutions (10^−1^ to 10^−6^) of MGM100 cells expressing the chaperonin plasmids were grown on LB medium supplemented with 0.5% glucose and 0.1 mM IPTG at 37°C and 45°C for 12 h. **(B)** ATPase assays of chaperonins. The ATPase activities of chaperonins GroEL and CPN60α/β1/β2 were measured in the absence and presence of functional co-chaperonin combinations at 25°C. **(C)** Refolding of denatured RrRubisco by chaperonins. Guanidine hydrochloride denatured RrRubisco (25 μM) was diluted 100-fold into refolding buffer containing 0.25 μM GroEL or CPN60α/β1/β2. The refolding reaction was initiated by the addition of 1 µM of the various co-chaperonins and 2 mM ATP, allowed to proceed for 1 h at 25 °C, and then stopped by the addition of 10 mM glucose and 2.5 U of hexokinase. RrRubisco carboxylation activity was measured by an assay based on ^14^C isotope labeling. The activity of native RrRubisco (0.25 μM) was set to 100%.

To test the propensity of the co-chaperonins to form homo-and hetero-oligomers, we co-expressed three co-chaperonin genes at the combinations shown in Fig. S3A in *E. coli.* The recombinant protein oligomers were purified and analyzed by SDS-PAGE and non-denaturing (ND)-PAGE. CPN20 alone and in combination with the other co-chaperonins formed stable hetero-oligomeric complexes *in vitro* (Fig. S3A). We then performed gel filtration to investigate the interaction between GroEL and CPN60αβ1β2 and the chloroplast co-chaperonins *in vitro.* These analyses revealed that CPN20, CPN11/20, CPN11/23, CPN20/23 and CPN11/20/23 all co-migrated with GroEL and CPN60αβ1β2 in high molecular mass complexes in the presence of ATP (Fig. S3B), suggesting the formation of common complexes. Subsequent ATPase assays showed that GroES, CPN11/20, and CPN11/20/23 inhibited the ATPase activities of GroEL and CPN60αβ1β2 (Fig. 2B), suggesting their functional interactions (Todd et al., 1993). Although CPN20/23 did not inhibit the ATPase activity of GroEL, it did inhibit that of CPN60αβ1β2 (Fig. 2B). Moreover, GroES and CPN11/20/23 were the most effective co-chaperonins in supporting the refolding of *Rhodospirillum rubrum* Rubisco (RrRubisco) by GroEL and CPN60αβ1β2 (Fig. 2C). In summary, the co-chaperonin combination CPN11/20/23 possessed superior activity in GroEL-and CPN60αβ1β2-mediated chaperonin function both *in vitro* and in *E. coli*, implying that CPN11/20/23 hetero-oligomers may occur as a general co-chaperonin in the chloroplast.

### The stoichiometry of CPN60 subunits in native CPN60αβ1β2 complexes

It has been discussed controversially whether the heptameric stacked rings of chloroplast chaperonins consist of homo-oligomeric α-and β-rings, or whether both rings are hetero-oligomeric (Dickson et al., 2000; Vitlin Gruber et al., 2013). To address this question, we purified endogenous CPN60αβ1β2 from *Chlamydomonas reinhardtii* chloroplasts. Using a combination of chloroplast isolation, ammonium sulfate precipitation and a series of chromatographies, we eventually obtained a preparation of endogenous CPN60αβ1β2 (eCPN60αβ1β2) that according to SDS-PAGE and ND-PAGE analyses was highly pure (Fig. 3A). eCPN60αβ1β2 migrated in ND-PAGE with exactly the same molecular mass as recombinant CPN60αβ1β2 (rCPN60αβ1β2), indicating that purified eCPN60α/β1/β2 forms a stable complex (Fig. 3A). However, the ATPase activity of eCPN60αβ1β2 was almost two times higher than that of rCPN60αβ1β2 which might be due to different stoichiometries of CPN60α/β1/β2 subunits within the complexes, or trace substrate contaminations in the purified eCPN60αβ1β2 (Fig. 3B). The addition of recombinant co-chaperonins CPN11/20/23 led to a significant inhibition of eCPN60αβ1β2 ATPase activity, indicating a functional interaction between CPN11/20/23 and eCPN60αβ1β2.

**Figure 3.**
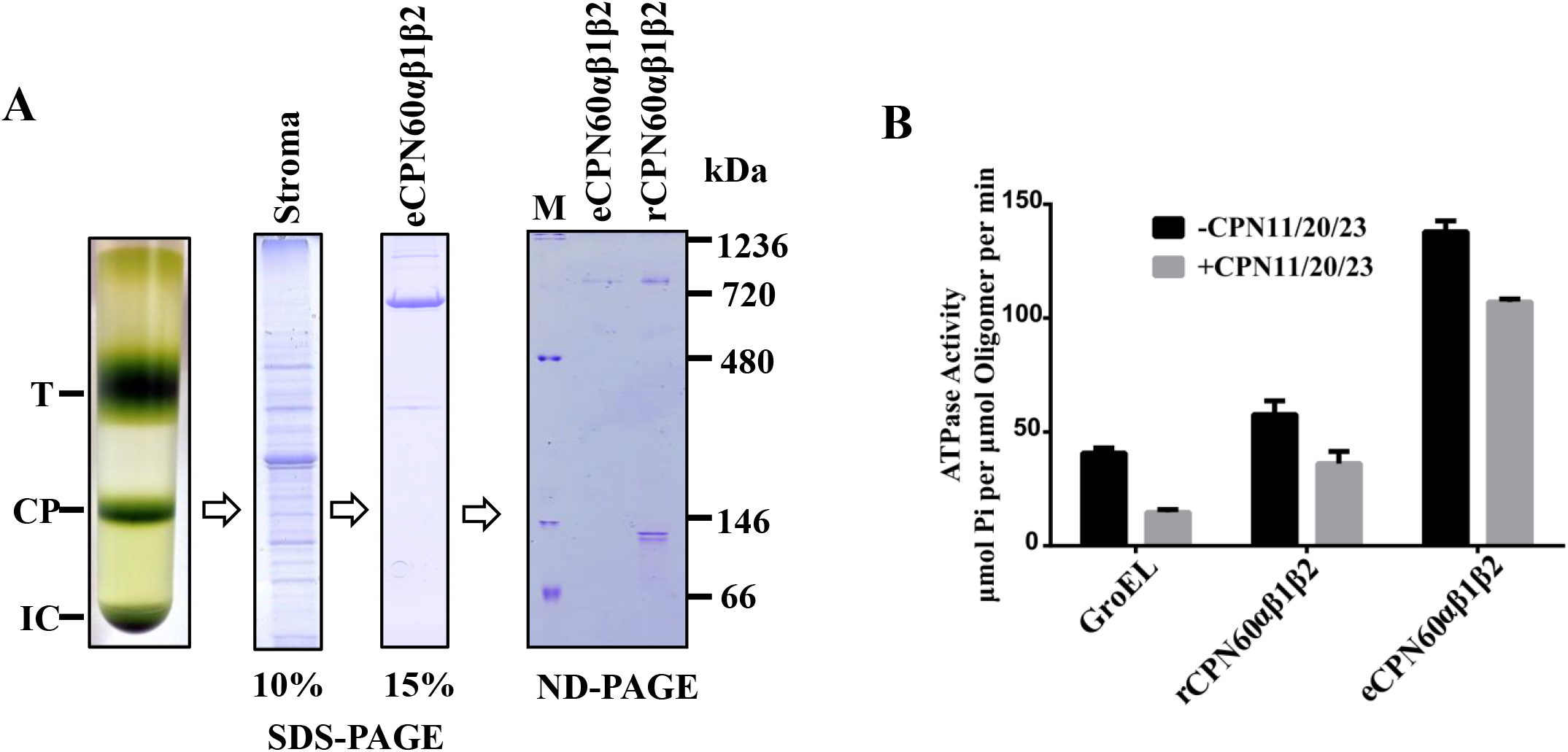
Purification of endogenous CPN60α/β1/β2 from *Chlamydomonas reinhardtii* chloroplasts. **(A)** Chloroplasts were separated by Percoll gradient centrifugation yielding thylakoids (T), chloroplasts (CP) and intact cells (IC). Chloroplasts were collected and osmotically lysed to obtain stroma protein. Endogenous chloroplast chaperonin complexes (eCPN60αβ1β2) were purified through ammonium sulfate precipitation, ion exchange chromatography, hydrophobic interaction chromatography and size exclusion chromatography. Purity and assembly state of eCPN60αβ1β2 were compared with the recombinant complex (rCPN60αβ1β2) by SDS-PAGE and ND-PAGE. **(B)** ATPase activity of native and recombinant chaperonins. The ATPase activity of eCPN60αβ1β2 was measured in the absence and presence of co-chaperonins CPN11/20/23 at 25°C by a NADH-coupled reaction. ATPase activities of GroEL and rCPN60αβ1β2 were measured for comparison.

To quantify the molar ratio of CPN60α/β1/β2 subunits within the eCPN60αβ1β2 hetero-oligomers, we employed a QconCAT protein harboring concatenated proteotypic (Q-)peptides that cover the three subunits of *Chlamydomonas* chloroplast chaperonins (Bai et al., 2015). The QconCAT protein was digested tryptically and analyzed by LC-MS/MS to obtain extracted ion chromatograms (XICs) for the Q-peptides (Fig. S4). These XICs allowed us to determine the ratios of the ion intensities of the Q-peptides that are released in equimolar amounts from the QconCAT protein. Gel bands corresponding to the denatured CPN60 monomers in SDS-polyacrylamide gels (from toal cell lysate protein, stroma lysate protein, purified endogenous eCPN60αβ1β2 and recombinantly expressed rCPN60αβ1β2) and to the native CPN60α/β1/β2 complex in ND-polyacrylamide gels (purified endogenous eCPN60αβ1β2 and recombinantly expressed rCPN60αβ1β2) were excised, subjected to in-gel tryptic digestion, and analyzed by LC-MS/MS to obtain XICs for the native peptides with sister Q-peptides in the QconCAT protein. Based on the XICs obtained for the native peptides and the information on the ratios of ion intensities obtained for the sister Q-peptides, we determined the stoichiometries of the endogenous CPN60 (eCPN60αβ1β2) was about 5 : 3: 6 or 6 : 2 : 6 for CPN60α : CPN60β1 : CPN60β2 (Table 1). It is unclear if the molar ration of each subunit in the native CPN60 complex is fixed, if so, we suspect that the two different stoichiometries results from the experimental procedures. Since the standard deviation in 6 : 2 : 6 stoichiometry is much smaller, we tend to think this ratio is real. Moreover, the subunit molar ratio of the recombinantly expressed CPN60 (rCPN60αβ1β2) was determined as 4 : 7: 3 for CPN60α : CPN60β1 : CPN60β2 (Table 1). This stoichometry is different from that of eCPN60 (6 : 2 : 6 for α : β1 : β2) and that of earlier report from our lab (5 : 6: 3 for CPN60α : CPN60β1 : CPN60β2)(Bai et al., 2015). The difference might result from the expression amount variations of each subunit in Chlamy and *E.coli* (the polycistron was used to express three subunits in *E.coli* BL21 in Bai’s paper, and each subunit was expressed in *E.coli* MGM100 with its own promotor and terminator in this work). We could not exclude the possibility that the recombinantly purified rCPN60 is the population of oligomers with varied subunits stoichiometry. The stoichiometry difference might also resulte from the experimental procedure as that of the native eCPN60 in this work.

**Table 1.** Stoichiometries of CPN60 subunits within CPN60α/β1/β2 complex.

| | Total protein | Stroma protein | eCPN60α/β1/β2 | | rCPN60α/β1/β2 | |
| --- | --- | --- | --- | --- | --- | --- |
| Subunit | Number (SDS-PAGE) | Number (SDS-PAGE) | Number (SDS-PAGE) | Number (ND-PAGE) | Number (SDS-PAGE) | Number (ND-PAGE) |
| CPN60α | 5.54±0.08 | 4.85±0.65 | 5.00±0.53 | 5.60±0.11 | 4.06±0.12 | 4.24±0.06 |
| CPN60β1 | 1.98±0.10 | 3.47±0.37 | 2.83±0.81 | 2.34±0.038 | 7.36±0.24 | 7.18±0.12 |
| CPN60β2 | 6.47±0.09 | 5.68±0.32 | 6.20±0.30 | 6.06±0.090 | 2.58±0.15 | 2.58±0.08 |

### Cryo-EM structure of eCPN60αβ1β2 and eCPN60αβ1β2-CPN11/20/23

We previously reported the crystal structure of the *Chlamydomonas* CPN60β1 homo-oligomer, which shares a similar structural topology with group I chaperonins (Zhang et al., 2016a). However, compared to the homo-oligomeric GroEL and Hsp60, the hetero-oligomeric chloroplast chaperonin system suggests an asymmetric structural organization which might be related to its unique function. To investigate the hetero-oligomeric structure of the chloroplast chaperonin and develop a detailed picture of its interaction with the chloroplast co-chaperonins, we employed single particle cryo-EM. We chose to determine two structures along the ATP-driven functional cycle: the first is the ADP-bound state of eCPN60αβ1β2, as obtained by incubating purified native CPN60αβ1β2 with ADP. The second is the folding-active state of eCPN60αβ1β2 formed with chloroplast co-chaperonins CPN11/20/23 in the presence of ADP. The sample was prepared by mixing purified eCPN60αβ1β2 and recombinant CPN11/20/23 at a molar ratio of ~1:1.5 in the presence of excessive amount of ADP. The cryo-EM images show that most of the particles reveal a homogeneous cylindrical shape for CPN60αβ1β2 and a bullet-shap for CPN11/20/23-CPN60αβ1β2 (Fig. S5).

Our reference-free two-dimensional (2D) class averages reveal multiple orientations and great details for both CPN60αβ1β2 and CPN11/20/23-CPN60αβ1β2 (Fig. S5C). Since the chloroplast chaperonin system is composed of different types of chaperonin and co-chaperonin subunits, no symmetry was imposed in the three-dimensional (3D) classification and refinement procedures (Fig. S6A). Two cryo-EM density maps were determined with overall resolutions of 4.06 Å and 3.82 Å for CPN60αβ1β2 and CPN11/20/23-CPN60αβ1β2, respectively, based on the gold-standard Fourier shell correlation(van Heel and Schatz, 2005) (Fig. S6B). The overall structures of CPN60αβ1β2 and CPN11/20/23-CPN60αβ1β2 both preserve the canonical architectures of GroEL and GroES-GroEL, respectively, despite their complex compositions (Fig. 4). The CPN60 subunit can also be divided into equatorial domain, intermediate domain and apical domain, while the CPN11/20/23 co-chaperonin complex consists of seven GroES-domain-like β-barrel structures with weak density of mobile loop regions reflecting the disordered nature of these loops. The domain rearrangements taking place during the transition of CPN60αβ1β2 to the co-chaperonin-bound folding-active state are also conserved among group I chaperonins, as indicated by the GroEL and GroEL-GroES models that were fitted into the density maps (Fig. S7). For both maps, the resolutions are not uniform: in general, the resolution in the apical domain is lower than that in the equatorial and intermediate domains (Fig. S7). It is noteworthy that the *trans*-ring of CPN11/20/23-CPN60αβ1β2, especially the apical and intermediate domain regions, was less well resolved, suggesting intrinsic flexibility or conformational heterogeneity of this hetero-oligomeric ring (Fig. S7). Since ATP and substrate bind to the open trans-ring in the GroEL-GroES recycling paradigm, the flexible nature of the *trans*-ring in CPN11/20/23-CPN60αβ1β2 may be beneficial for its binding with ATP and substrate.

**Figure 4.**
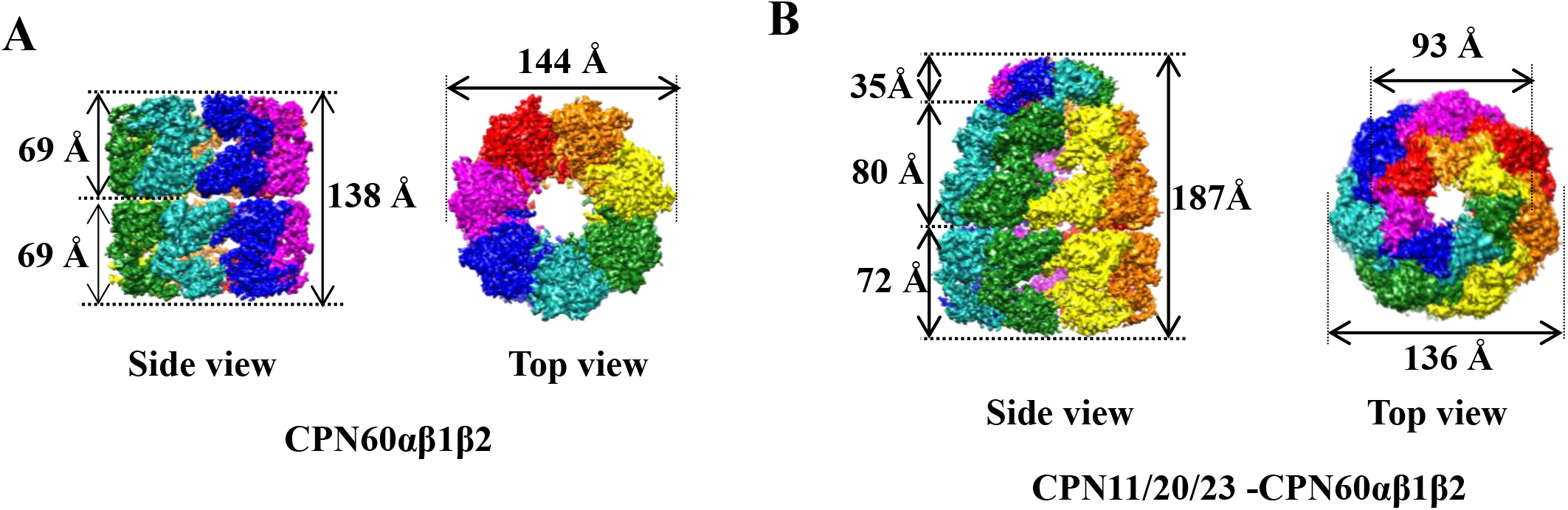
Symmetry-free cryo-EM maps of chloroplast chaperonins. **(A)** Top and side views of CPN60αβ1β2. **(B)** Top and side views of CPN11/20/23-CPN60αβ1β2. To distinguish between the seven chaperonin and co-chaperonin subunits they are shown in different colors.

### Conformational heterogeneity of chaperonin subunits

The chloroplast chaperonin system consists of multiple homologous co-chaperonin and chaperonin subunits. Since the density maps were calculated without imposing any symmetry, we attempted to identify the CPN60α-, β1-, and β2-subunits within the cis-ring of the relatively better resolved CPN11/20/23-CPN60αβ1β2 complex. The extremely high sequence similarity among the three CPN60 subunits (Fig. S8A) and lacking of adequate length of insertions result in impossibility to unambiguously distinguish the three subunits of CPN60 at the current resolution although the side-chain densities are visible in most portions of the equatorial domain and GroEL could be clearly distinguished from CPN60 based on prominent large side-chain densities (Fig. S9).

Group II chaperonin TRiC/CCT consists of eight distinct subunits that differ from each other by their ability to recognize substrates and their allosteric cooperativity along the ATPase cycle (Joachimiak et al., 2014; Lopez et al., 2015; Reissmann et al., 2012; Zang et al., 2016). We wondered whether a similar diversification might also exist in the CPN60αβ1β2 complex. However, the rotational correlation analysis of the upper ring of CPN60αβ1β2 as well as the *cis*-ring of CPN11/20/23-CPN60αβ1β2 (Figs. 5A and 5B) reveal an obvious pseudo 7-fold symmetry of the rings in both conformational states, instead of obvious conformational heterogeneity among the intra-ring subunits. In addition, we also calculated the rotational correlation between the two heterogeneous heptameric rings of CPN60αβ1β2, revealing high conformational similarities between the two rings as well as among the seven intra-ring subunits (Fig. 5B).

**Figure 5.**
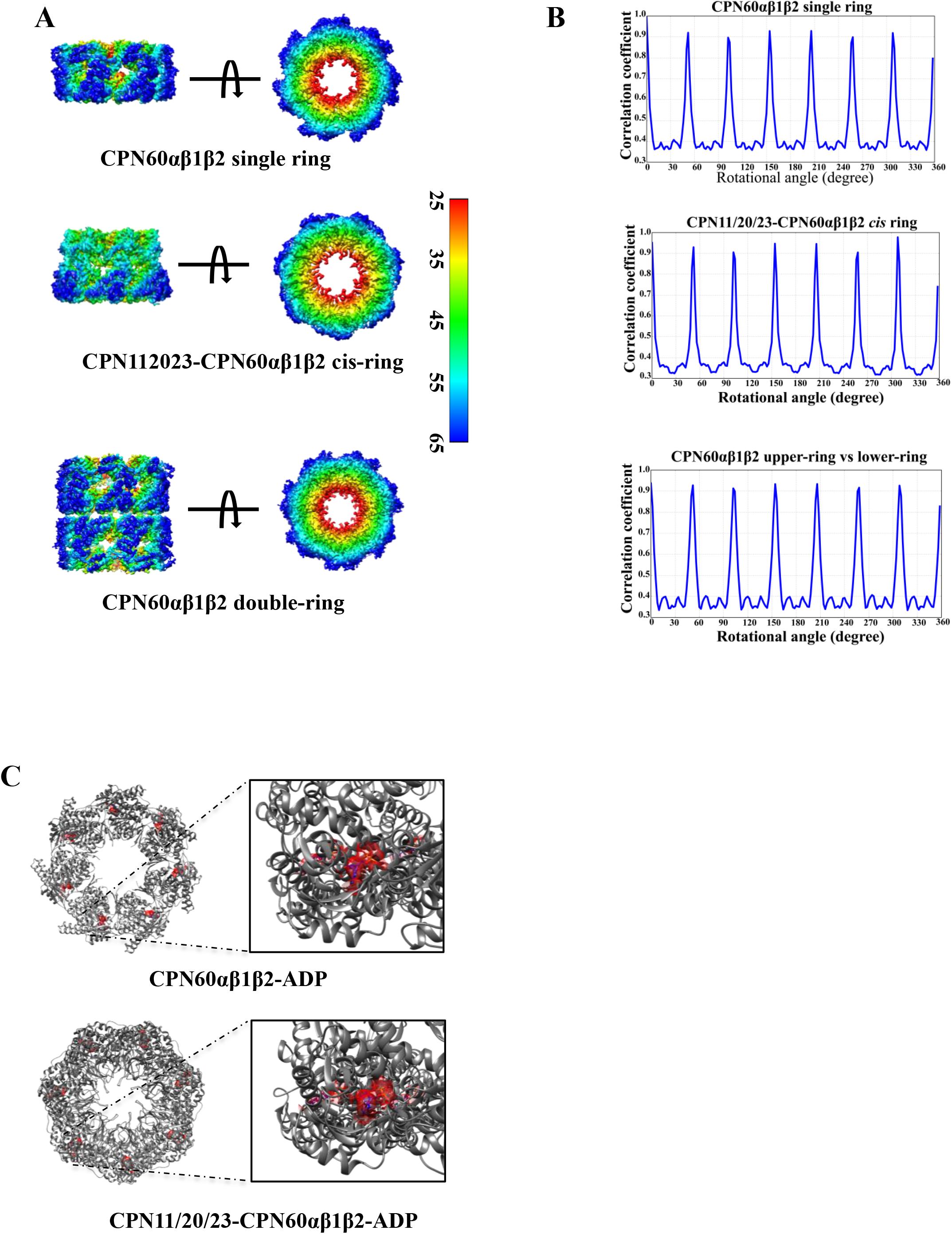
Comparison of chaperonin subunits in two ATPase cycle conformations. **(A)** Side and top views of CPN60αβ1β2 upper-ring, CPN11/20/23-CPNαβ1β2 *cis*-ring, and the intact double ring structure of CPN60αβ1β2. Maps are colored according to their cylinder radius (unit: Å). **(B)** Cross-correlation coefficients between the map of the CPN60αβ1β2 upper-ring and the CPN11/20/23-CPNαβ1β2 *cis*-ring and their symmetric references and cross-correlation coefficients between the map of CPN60αβ1β2 upper-ring and lower-ring. The plots indicate that the two hetero-oligomeric rings of CPN60αβ1β2 upper-ring and CPN11/20/23-CPNαβ1β2 *cis*-ring show nearly perfect C7 symmetry and the two rings of CPN60αβ1β2 shows D1 symmetry. **(C)** Localization of the ADP density (red) in the difference maps made between the CPN60αβ1β2 upper-ring and the CPN11/20/23-CPNαβ1β2 *cis*-ring. The optimized GroEL and GroES-GroEL models were taken as a framework to show the location of ADP. The atomic structure of ADP is fitted into the density and shown in the zoom-in picture.

The protein folding cycle of chaperonins is driven by nucleotide binding and hydrolysis. To investigate whether all CPN60α/β1/β2 subunits bind to nucleotide, we inspected the nucleotide-binding pockets of all the seven subunits in the CPN60αβ1β2 upper-ring and in the CPN11/20/23-CPN60αβ1β2 *cis*-ring, and observed that, in both of the these rings, all the seven nucleotide-binding pockets were occupied by ADP in the maps of both ring types (Fig. 5C). It has been reported that the ATP-binding affinities of different TRiC/CCT subunits vary, which results in a staggered induction of conformational changes in TRiC subunits upon ATP-binding (Reissmann et al., 2012; Zang et al., 2016). To test the possibility that CPN60α/β1/β2 subunits vary in their ATP-binding affinity, the surface electrostatic potential of the nucleotide-binding pockets was calculated based on atomic models fitting into the same CPN60 subunit electron density of the CPN11/20/23-CPN60αβ1β2 map. Notably, the electrostatic potential surfaces of the CPN60α/β1/β2 nucleotide-binding pockets are similar, but that of CPN60α is more negatively charged, implying a lower affinity for nucleotides (Fig. S10).

The physical environment of the central cavity of GroEL was proposed to be important for its ability to assist in protein folding. Therefore, we computed the electrostatic potential and amino acid hydrophobicity of individual CPN60α/β1/β2 subunits and compared them with GroEL. The overall distribution of hyrophobic amino acids is similar among CPN60 subunits, except for some hydrophobic proportions lining the surface of the apical and equatorial domains of CPN60α (Fig. 6A). Regarding the electrostatic potentials, are overall similar among the subunits, but CPN60β1 and CPN60β2 are more positively charged comparied with that of CPN60α (Fig. 6B). Therefore, while the amino acid hydrophobicity and electrostatic potential distribution in the inner chamber of GroEL are symmetrical, they are asymmetrical in CPN60αβ1β2 complex, which might be important. Therefore, although the different CPN60 subunits have little structural differences in CPN60αβ1β2 and CPN11/20/23-CPN60αβ1β2, their physicochemical properties are different. The heterogeneous nature of CPN60αβ1β2, particularly regarding the surface property of its protein folding nano-cage, might be crucial for the folding of specific obligate chloroplast substrates.

**Figure 6.**
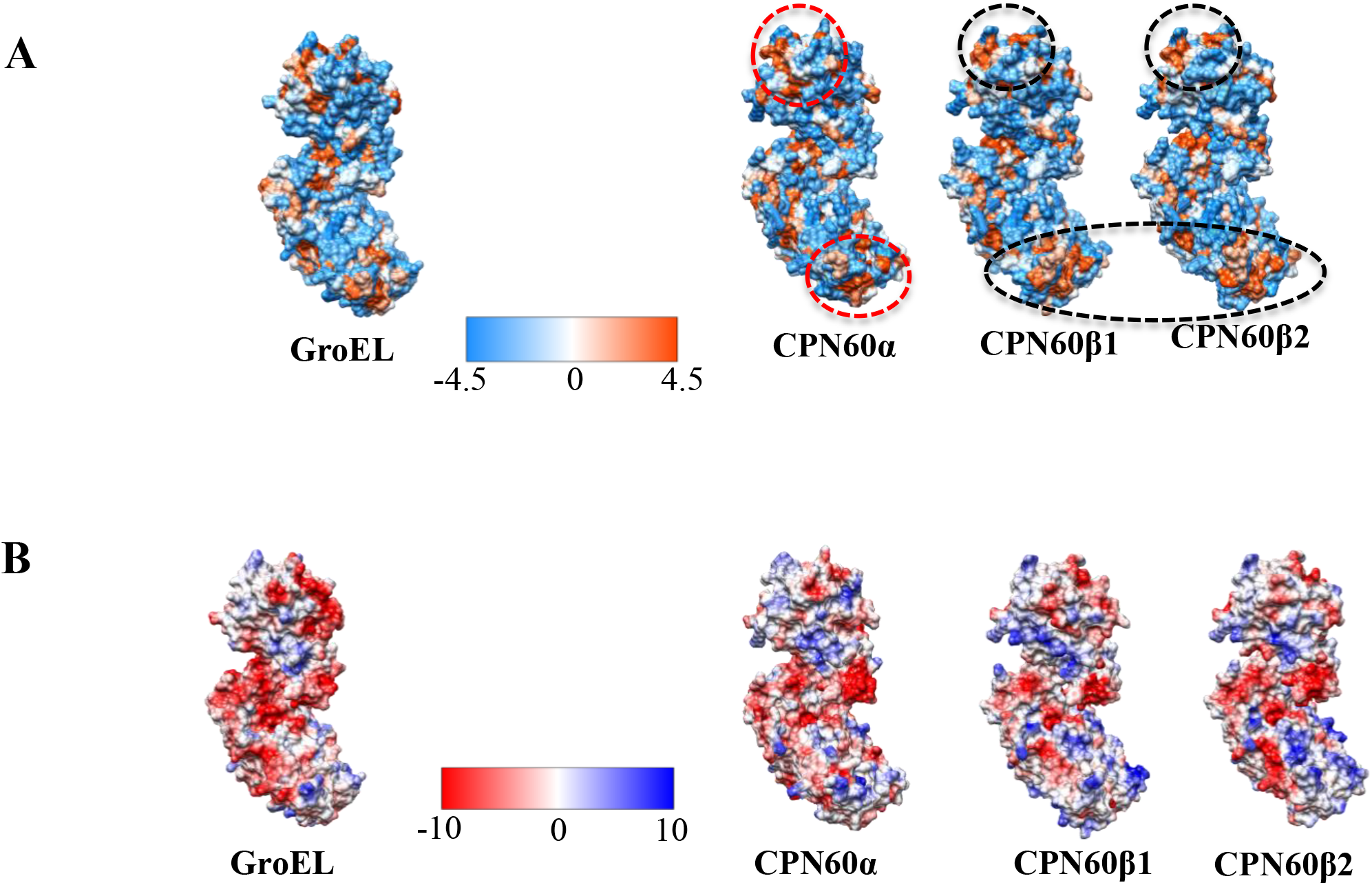
Surface properties of CPN60 subunits towards central cavity side. **(A)** Individual subunits of GroES-GroEL and CPN11/20/23-CPN60αβ1β2 with central cavity surface and side-chain properties are shown: hydrophilic (sky blue), hydrophobic (orange), and main chain (white). **(B)** The central cavity surface electrostatic potentials of individual subunits from GroES-GroEL and CPN11/20/23-CPN60αβ1β2 was calculated with UCSF Chimera’s Coulombic Surface Coloring module with red standing for a negative potential, white for near neutral, and blue for a positive potential.

### Structure of the chloroplast co-chaperonin and its interaction with the chaperonin

By regulating the inner chamber volume or its chemico-physical surface properties, co-chaperonins could endow the chaperonin system with features that enable them to assist in the folding of specific substrates. As we could show that the chloroplast CPN11/20/23 co-chaperonin complex functionally cooperates with both CPN60αβ1β2 and GroEL (Figs. 1 and 2), we decided to examine its structure. Although the resolution was relatively low in the CPN11/20/23 region of the CPN11/20/23-CPN60αβ1β2 map, we could clearly distinguish seven GroES-like domains (Fig. 7A). Rotational correlation analysis of CPN11/20/23 revealed high similarity among the GroES-like domains of the different co-chaperonin subunits (Figs. 7A and 7B). Hence, the CPN11/20/23 complex matches well with the heptameric *cis*-ring of CPN60α/β1/β2, therefore solving the symmetry problem by co-chaperonin subunit hetero-oligomerization (Guo et al., 2015; Tsai et al., 2012; Vitlin Gruber et al., 2014; Weiss et al., 2009).

**Figure 7.**
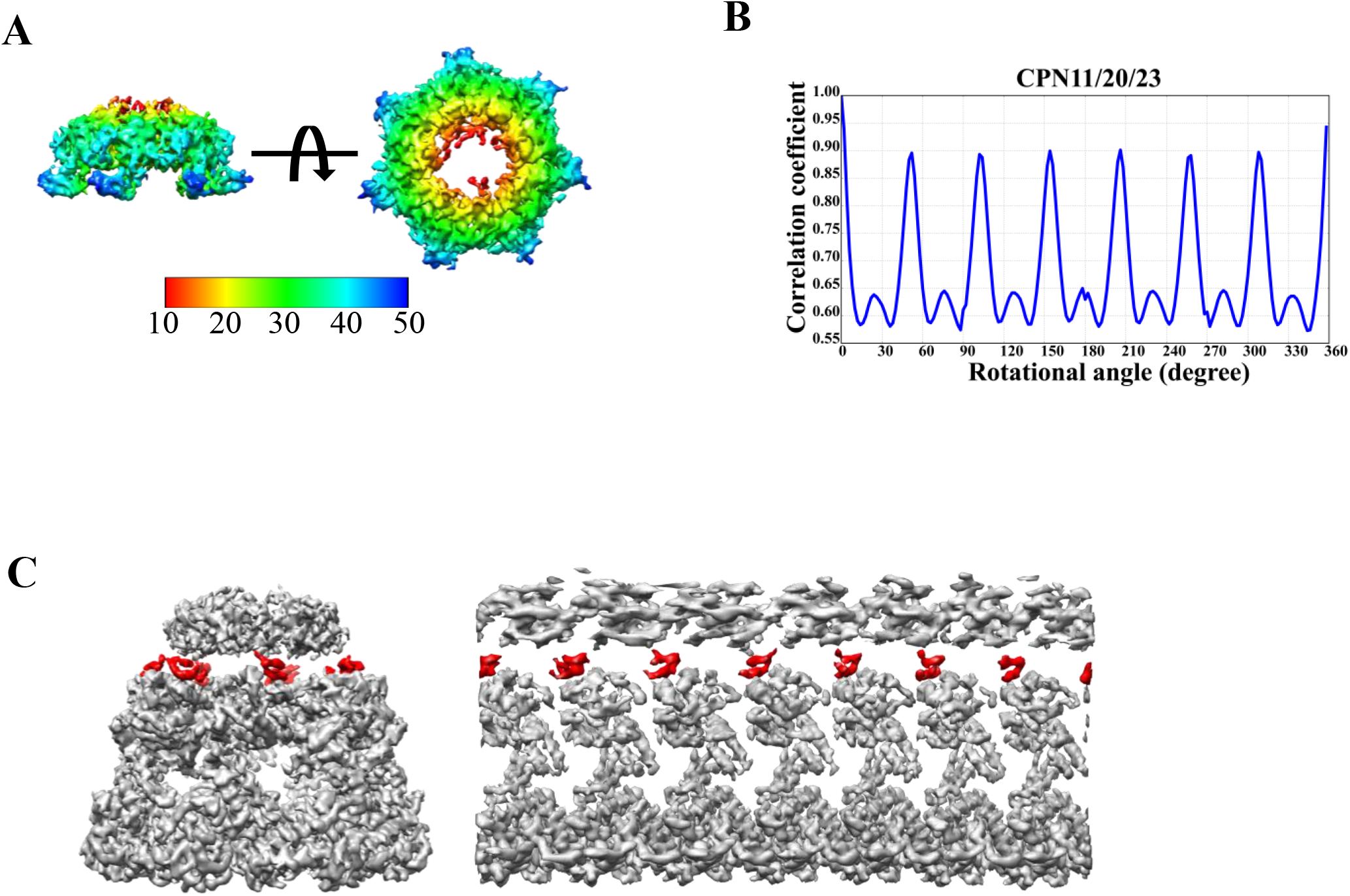
Cryo-EM structure of the CPN11/20/23 co-chaperonin complex. **(A)** Side and top views of the CPN11/20/23 map. The map is colored according to its cylinder radius (unit: Å). The loop structures in the dome region show an asymmetrical distribution. **(B)** Coefficients of cross-correlations between the map of CPN11/20/23 and its symmetrical reference. The peaks in each curve are nearly identical, indicating that the 10-kDa domain of CPN11/20/23 also adopts psudeo-C7 symmetry. **(C)** CPN11/20/23 interacts with all seven subunits of CPN60αβ1β2.The map (left) and unwrapped map (right) of the *cis*-ring of CPN11/20/23-CPNαβ1β2 are shown with the mobile loop density mediating the interaction between CPN11/20/23 and CPNαβ1β2 colored in red.

The densities of the flexible mobile loop regions, mediating the interaction of chaperonin with co-chaperonin, are relative weak in the structure (Fig. 7C). However, the β-hairpin loop structures of the co-chaperonins that directly bind to the chaperonin subunits are clearly visible, presumably because of their stability (Fig. 7C). It has been shown previously that the apical domains of CPN60α and CPN60β subunits have diverged with respect to their recognition of GroES (Zhang et al., 2016b). As shown by the unwrapped density map, all CPN60 subunits interact with GroES-like domains in the CPN11/20/23 complex (Fig. 7C). Sequence alignments revealed that the N-terminal GroES-like domain of CPN20 lacks the amino acids corresponding to a β-hairpin that forms the roof of the dome-shaped co-chaperonin (Fig. S8B). Accordingly, in the CPN11/20/23 map we observed that the dome structure was not uniformly distributed in the seven GroES-like domains (Fig. 7A), indicating that different combinations of co-chaperonin subunits may result in diverse cavity roof properties that might be required for assisting in the folding of specific substrate proteins.

## Discussion

The chloroplast chaperonin was first recognized as a protein that transiently binds Rubisco large subunits before they assemble into the holoenzyme (Barraclough and Ellis, 1980; Ellis et al., 1989; Hemmingsen and Ellis, 1986). Two features set the chloroplast chaperonin apart from the prototypical GroEL/ES system in *Escherichia coli*: (1) its irreplaceable ability to assist in the folding and assembly of the Rubisco holoenzyme; (2) the hetero-oligomeric nature of chloroplast chaperonin and co-chaperonins, consisting of Cpn60α-and Cpn60β-subunits and co-chaperonin Cpn10 and Cpn20-subunits, respectively (Aigner et al., 2017; Roy, 1989; Vitlin Gruber et al., 2013; Zhao and Liu, 2017). In a previous study the recombinant chloroplast co-chaperonin subunits CPN11, CPN20 and CPN23 from *Chlamydomonas* form various hetero-oligomeric complexes when mixed *in vitro* (Tsai et al., 2012). Here we provide evidence for the existence of chloroplast co-chaperonin hetero-oligomers in *Chlamydomonas* also *in vivo,* as CPN11, CPN20 and CPN23 could be co-immunoprecipitated with CPN60αβ1β2 from chloroplast stroma (Fig. 1). We took advantage of the GroES/EL deficient strain MGM100 to systematically study the functionality of *Chlamydomonas* co-chaperonin subunit combinations with GroEL or the native partner, CPN60αβ1β2. We showed that CPN20 alone and all possible combinations of chloroplast co-chaperonin subunits can perform co-chaperonin function in *E. coli,* with both GroEL and CPN60αβ1β2. However, with GroEL all chloroplast co-chaperonins failed to fully complement MGM100 under heat stress conditions, implying the requirement for a species-specific cooperation to cope with unfolded proteins accumulating under heat stress. An intriguing phenomenon was that CPN20-GroEL was able to complement MGM100 under ambient conditions, but unable to assist in the folding of denatured RrRubisco *in vitro* (Fig. 2), consistent with results from a previous study (Tsai et al., 2012). This suggests that CPN20-GroEL assists in the folding of proteins essential for cell survival, but not of any substrate. Several biochemical assays (Figs. 2, S2 and S3), including co-migration in gel filtration, ATPase activity inhibition, and *in vitro* refolding of RrRubisco, indicated that different chloroplast co-chaperonin compositions had different bioactivities, which may correlate with different functions *in vivo*. In all our biochemical assays, the co-chaperonins (GroES or CPN20 combinations) were functional with both bacterial and green algal chaperonin, but the homologous CPN11/20/23-CPN60αβ1β2 complex was more capable (Fig. 2A), similar to the chloroplast and bacterial Hsp70 system (Dorn et al., 2010). These results indicate a strong functional conservation between the bacterial and green algal chaperonin systems despite the substantial subunit differentiation. However, the homo-oligomeric CPN20, which failed to inhibit GroEL ATPase activity and assist in RrRubisco folding *in vitro*, showed normal co-chaperonin activity when cooperating with CPN60αβ1β2, to the extent that CPN20-CPN60αβ1β2 could even complement MGM100 under heat stress conditions, indicating the high specificity of CPN20 in the chloroplast chaperonin system (Figs. 2 and S2). Our results support the idea that the chloroplast co-chaperonin complex composed of diversified subunits may endow the chaperonin system to preferentially assist in the folding of specific substrate proteins.

GroES/GroEL structures in different conformational states have been studied extensively in the past and have provided invaluable insights into the working mechanism of group I chaperonins as protein folding machines (Boisvert et al., 1996; Chen et al., 2013; Clare et al., 2012; Ranson et al., 2006; Xu et al., 1997). A structure of the human homo-oligomeric mitochondrial chaperonin has also been determined by X-ray crystallography and revealed a symmetrical football shape (Nisemblat et al., 2015). Recently, we have reported the crystal structure of the homo-oligomeric CPN60β1 chloroplast chaperonin from *Chlamydomonas,* which shares a similar topology with GroEL (Zhang et al., 2016a). However, a structure of the authentic hetero-oligomeric chloroplast chaperonin even after many years of effort is still elusive, presumably due to the labile nature of the recombinant hetero-oligomeric complex. It has been proposed that the chloroplast chaperonin consists of two homo-oligomeric rings, each composed of α-or β-subunits (Dickson et al., 2000; Vitlin Gruber et al., 2013). Here we present cryo-EM structures of the endogenous CPN60αβ1β2 in ADP binding state at 4.06 Å resolution, and in the co-chaperonin CPN11/20/23 and ADP binding state at 3.82 Å resolution. Our data are in disfavor of this idea, as the stoichiometry of α:β1:β2 in the native chaperonin is ~6:2:6 (table 1) and the rotational analysis between the two rings of CPN60αβ1β2 also showed internal symmetry (Fig. 5). Although, the cryo-EM structure at current resolution did not allow us to distinguish α-and β-subunits within the ring, it is sufficient to reveal a nearly perfect pseudo seven-fold symmetry of the seven chaperonin subunits within the ring in the two conformational states by rotational correlation analysis (Fig. 5). This is different from the group II chaperonin TRiC, which consists of eight distinct subunits within each ring which diverge in substrate binding sites and activities (Cong et al., 2012; Joachimiak et al., 2014; Leitner et al., 2012; Reissmann et al., 2012). This may be attributed to the higher sequence homology of chloroplast chaperonin subunits when compared with TRiC subunits. Moreover, CPN60αβ1β2-ADP and CPN11/20/23-CPN60αβ1β2-ADP are start and end points along the ATP-driven chloroplast chaperonin reaction cycle, *i.e.,* varied allosteric states of subunits during folding may not be observed in our structures. It is noteworthy that the trans-ring of CPN11/20/23-CPN60αβ1β2 was less well resolved compared with the cis-ring (Fig. S7), which was not seen in any of the GroES/GroEL structures, indicating a dynamic feature of the CPN60 subunits in the trans-ring. There is a paradox that the apical domains of the chaperonin simultaneously mediate the binding of the co-chaperonin and the substrate protein. A cryo-EM structure of GroEL-GroES encapsulating a substrate protein showed that the GroEL cis-ring apical domains can simultaneously bind GroES and substrate protein by breaking the seven-fold symmetric pattern (Chen et al., 2013). Therefore, it is tempting to speculate that the chloroplast chaperonin system might solve this paradox through subunit differentiation. As we have reported previously, Cpn60α/β showed different binding affinity for co-chaperonin and substrate protein (Zhang et al., 2016b). To address this question, future structural study on CPN11/20/23-CPN60α/β1/β2 with bound substrate protein will be required. Studies on TRiC/CCT revealed the eight paralogous subunits to generate an asymmetric power stroke driving the chaperonin TRiC/CCT folding cycle (Reissmann et al., 2012; Zang et al., 2016). The ATP binding pockets of CPN60α/β1/β2 based on the pseudo-atomic model showed slight differences in surface properties, suggesting a similar mechanism may exist also in the chloroplast chaperonin system (Fig. S10).

The structure of the CPN11/20/23 chloroplast co-chaperonin, consisting of one Cpn10 and two Cpn20 subunits, also showed a pseudo seven-fold symmetry (Fig. 7). This also substantiates earlier proposal that chloroplast co-chaperonins solve the symmetry mismatch problem through hetero-oligomerization (Tsai et al., 2012; Weiss et al., 2009). In addition, the very similar structure between CPN11/20/23 and GroES may explain our biochemical data that both co-chaperonins support GroEL activity to a similar extent (Figs. 1 and 2). Sequence alignments of GroES and *Chlamydomonas* chloroplast co-chaperonin subunits revealed that the N-terminal GroES-like domain of CPN20 lacks the sequence which forms a roof-like β-hairpin in the dome-shaped co-chaperonin complex, while CPN10 and both GroES-like domains of CPN23 retain this sequence (Fig. S8B). The different combinations of the co-chaperonin subunits varying in the roof-like sequences may result in diverse properties the roof formed by different chloroplast co-chaperonin complexes. This idea is supported by our symmetry-free CPN11/20/23 cryo-EM map, which shows an asymmetric pattern of the roof-like loops (Fig. 7). Finally, *Chlamydomonas* CPN20 exhibits a species-specific co-chaperonin activity when cooperating with CPN60αβ1β2, and the interaction between CPN20 and CPN60αβ1β2 is an obligatory symmetry mismatch. Further structural studies will shed light on how the CPN20-CPN60αβ1β2 partnership works.

## Materials and Methods

### Plasmid construction for the co-expression of chaperonins and co-chaperonins

To construct a vector for the co-expression of multiple chaperonin and co-chaperonin subunits, the Ptac promoter and λt_0_ terminator of pQLinKN were replaced by the T7 promoter and terminator. For this, a fragment containing the LINK1 and LINK2 sequence was amplified by PCR with the primers 5’-AGTAACAACACCATTTAAATGGAGT-3’ and 5’-ACAATTGAATCTATTATAATTGTTA-3’ using plasmid pQLinkN as template. Then the fragment containing T7 promoter and terminator was amplified by PCR with the primers 5’-TATAATAGATTCAATTGTTAATACGACTCACTATAGGGGA-3’ and 5’-TTAAATGGTGTTGTTACTCAAAAAACCCCTCAAGACCC-3’ using plasmid pET11a as template. At last, the two fragments were ligated together by seamless cloning (Genebank Biosciences Inc) yielding plasmid pQLinkT. *groES* and *groEL* genes were ampilified by PCR from the GroES-pET11a and GroEL-pET11a plasmids (lab stocks) and individually cloned into pQLinkT at NdeI/BamHI sites. *CPN10*, *CPN20*, *CPN23*, *CPN60α*，*CPN60β1*，*CPN60β2* genes that encodes the mature chloroplast proteins were amplified by PCR on cDNA from *Chlamydomonas reinhardtii* CC400 and individually cloned into pQLinkT at NdeI/BamHI sites. A PacI-digested fragment from one pQLinkT plasmid was inserted into the SwaI site of another pQLinkT plasmid by ligation-independent cloning as described previously (Scheich et al., 2007). In this way, pQLinkT co-expression plasmids can accept any combination of chaperonin and co-chaperonin genes. All the co-expression plasmids used in this study were generated by this method. All clonings were confirmed by Sanger sequencing.

### Expression and purification of recombinant proteins

<u>GroEL:</u> *E. coli* BL21 (DE3), transformed with plasmid GroEL-pQLinkT, was grown in 4 liters of LB medium containing 100 μg/mL ampicillin at 37°C. GroEL expression was induced by the addition of 1 mM Isopropyl β-D-1-thiogalactopyranoside at an OD_600_ of ~0.5 and the cells were harvested 3 hours later. Unless stated otherwise, subsequent steps were performed at 4 °C. Cell pellets were resuspended in 120 ml lysis buffer (50 mM Tris-HCl pH 8.0, 30 mM NaCl, 1 mM DTT, 1 mM EDTA, 1 mM PMSF), and lysed using ultrasonic tissue homogenizers. The debris was removed by centrifugation (35,000g, 1 h) and the supernatant was passed through a 0.22 μm filter. Aliquots of the filtrate were applied to Source 30Q column pre-equilibrated with lysis buffer. Proteins were eluted using a NaCl gradient from 30 mM to 1 M in five column volumes. The GroEL-containing fractions were collected and concentrated using Amicon Ultra-15 Centrifugal Filter Units with nominal molecular weight limits (NMWL) of 100 kDa. Then the concentrated proteins were subjected to a Superdex 200 gel filtration column pre-equilibrated with buffer containing 50 mM Tris-HCl pH 7.5, 150 mM NaCl, 1 mM EDTA. Fractions containing purified GroEL were eluted from the void volumn. Purified GroEL was concentrated to ~50 mg/mL, supplemented with 5% glycerol, flash-frozen in liquid nitrogen, and stored at −80°C.

<u>CPN60αβ1β2:</u> recombinant CPN60αβ1β2 was expressed in the conditionally *groE* operon depleted *E. coli* strain MGM100 to avoid contaminations by GroEL. *E. coli* MGM100, transformed with the plasmid CPN60αβ1β2-pQLinkT, was grown in 4 liters of LB medium containing 100 μg/mL ampicillin, 100 μg/mL kanamycin, 1 mM diaminopimelic acid (DAP), 0.5 mM glucose at 37°C. CPN60αβ1β2 expression was induced with 0.1 mM Isopropyl β-D-1-thiogalactopyranoside at an OD_600_ of ~0.5 at 25°C and cells were harvested 8 hours later. Unless stated otherwise, subsequent steps were performed at 4 °C. Cell pellets were resuspended in 120 ml lysis buffer (50 mM Tris-HCl pH 8.0, 30 mM NaCl, 1 mM DTT, 1 mM EDTA, 1 mM PMSF) and further lysed using ultrasonic tissue homogenizers. The debris was removed by centrifugation (35000g, 1 h) and the supernatant was passed through a 0.22 μm filter. Aliquots of the filtrate were applied to a Source 30Q column pre-equilibrated with lysis buffer. Proteins were eluted using a NaCl gradient from 30 mM to 500 mM in five column volumes. CPN60αβ1β2 containing fractions were collected and (NH_4_)_2_SO_4_ was added to a final concentration of 500 mM. Then the above aliquots were applied to a Phenyl-sepharose column pre-equilibrated with 30 mM Tris-HCl pH 8.0, 500 mM (NH_4_)_2_SO_4_. Proteins were eluted using a (NH_4_)_2_SO_4_ gradient from 500 mM to 0 mM in five column volumes. The CPN60αβ1β2containing fractions were collected and concentrated using Amicon Ultra-15 Centrifugal Filter Units with nominal molecular weight limits (NMWL) of 100 kDa. Then the concentrated proteins were subjected to a Superdex 200 gel filtration column, pre-equilibrated with buffer containg 50 mM Tris-HCl pH 7.5, 150 mM NaCl, 1 mM EDTA. Fractions containing purified CPN60αβ1β2 were eluted from the void volume. Purified CPN60α/β1/β2 was concentrated to ~20 mg/mL, supplemented with 5% glycerol, flash-frozen in liquid nitrogen, and stored at −80°C.

<u>GroES, CPN20, CPN1120, CPN1123, CPN2023, CPN11/20/23:</u> all co-chaperonins were purified with the same method. *E. coli* BL21 (DE3) cells were transformed with plasmids GroES-pQLinkT, CPN20-pQLinkT, CPN11/20-pQLinkT, CPN11/23-pQLinkT, CPN20/23-pQLinkT, and CPN11/20/23-pQLinkT. Transformed single clones were inoculated in 4 liters of LB medium containing 100 μg/mL ampicillin and grown at 37°C. Protein expression was induced by the addition of 1 mM Isopropyl β-D-1-thiogalactopyranoside at an OD_600_ of ~0.5 and cells were harvested 3 hours later. Unless stated otherwise, subsequent steps were performed at 4 °C. Cell pellets were resuspended in 120 ml lysis buffer (50 mM Tris-HCl pH 8.0, 30 mM NaCl, 1 mM DTT, 1 mM EDTA, 1 mM PMSF) and lysed using ultrasonic tissue homogenizers. Debris was removed by centrifugation (35,000g, 1 h) and the supernatant was passed through a 0.22 μm filter. In the next step, aliquots of the filtrate were applied to a Source 30Q column pre-equilibrated with buffer containing 50 mM Tris-HCl pH 8.0, 10 mM NaCl, 1 mM DTT, 1 mM EDTA. The protein was eluted using a NaCl gradient from 10 mM to 500 mM in five column volumes. The co-chaperonin containing fractions were collected and concentrated using Amicon Ultra-15 Centrifugal Filter Units with nominal molecular weight limits (NMWL) of 10 kDa. Then the concentrated proteins were subjected to Superdex 75 gel filtration column pre-equilibrated with buffer containing 50 mM Tris-HCl pH 7.5, 150 mM NaCl, 1 mM EDTA. Purified co-chaperonins were concentrated to ~50 mg/mL, supplemented with 5% glycerol, flash-frozen in liquid nitrogen, and stored at −80°C.

<u>CPN11_6xHis_2023:</u> *E. coli* BL21 (DE3) cells, transformed with plasmids CPN11_6xHis_20/23-pQLinkT. Culturing, induction of protein expression and cell harvest were done as described for the other co-chaperonins. For affinity purification, cells were resuspended in lysis buffer (50 mM NaH_2_PO_4_, 300 mM NaCl and 10 mM imidazole) and lysed by ultra-sonication. Cell debris was removed by centrifugation (35,000g, 1 h) and the supernatant was passed through a 0.22 μm filter. Soluble proteins were applied slowly to a Ni-NTA gravity column. After washing with lysis buffer containing 25 mM imidazole, proteins were eluted with lysis buffer containing 200 mM imidazole. Eluted proteins were dialyzed into buffer containing 50 mM Tris-HCl pH 7.5, 150 mM NaCl, 1 mM EDTA. Purified CPN11_6xHis_2023 proteins were concentrated to ~50 mg/mL, supplemented with 5% glycerol, flash-frozen in liquid nitrogen, and stored at −80°C.

### Chloroplast isolation and stroma extraction

Chloroplast isolation from *Chlamydomonas reinhardtii* was performed as described previously with minor modifications (Mason et al., 2006; Teng et al., 2012). The cell-wall-deficient strain of *Chlamydomonas reinhardtii* CC-400 (*cw15 mt+*), obtained from the Chlamydomonas resource center (https://www.chlamycollection.org/), was inoculated into TAP medium and grown mixotrophically under continuous light (45 μmol photons m^−2^ s^−1^) at 20°C. After 2 days, the cell cultures were switched to conditions of synchronous lighting (12 h light/12 h dark) for two light/dark cycles. Cells were harvested at 4 h into the third light cycle and broken by a Dounce tissue grinder (WHETON). Whole cells and crude chloroplasts were then loaded onto a discontinuous Percoll gradient (20%:45%:65%), and centrifuged for 15 min at 4200 g. Intact chloroplasts were collected from the 45–65% Percoll interface and washed with isolation buffer (containg 300 mM D-sorbitol) to remove the Percoll. Washed chloroplasts were lysed osmotically by resuspension in ice-cold buffer containing 10 mM Tris-HCl pH 8.0, 1 mM EDTA for 1 h. Debris was removed by centrifugation at 25,000 g for 90 min. The supernatant stroma was collected, supplemented with 5% glycerol, flash-frozen in liquid nitrogen, and stored at −80°C until use.

### Purification of endogenous chloroplast chaperonin CPN60αβ1β2

Stroma protein was thawed on ice and (NH_4_)_2_SO_4_ was added very slowly to a final concentration of 1.5 M with a magnetic stirrer. Precipitated proteins were removed by centrifugation for 10 min at 25,000g. The supernatant fractions containing enriched CPN60αβ1β2s were diluted to 0.75 M (NH_4_)_2_SO_4_ with 30 mM Tris-HCl pH 8.0 and applied to a 5 ml HiTrap Phenyl HP column. The column was pre-equilibrated with 30 mM Tris-HCl pH 8.0, 1 mM EDTA and eluted with a linear gradient of 10 column volumes from 0.75 M to 0 M (NH_4_)_2_SO_4_. After hydrophobic chromatography, CPN60αβ1β2 containing fractions were diluted to a lower conductivity (less than 10 mS/cm using 30 mM Tris-HCl pH 8.0, 1 mM EDTA) and applied to a 5-mL HiTrap Q HP column. The column was pre-equilibrated with 30 mM Tris-HCl pH 8.0, 50 mM NaCl, 1mM EDTA and developed with a linear gradient of 15 column volumes from 1 M to 50 mM NaCl. Fractions containing CPN60αβ1β2 were checked by immunoblotting. In order to separate CPN60αβ1β2 from Rubisco holoenzyme, the eluted fractions containing CPN60αβ1β2 were further fractionated on a MonoQ 5/50 GL column pre-equilibrated with 30 mM Tris-HCl pH 8.0, 50 mM NaCl, 1mM EDTA. The column was also developed with a linear gradient of 15 column volumes from 1 M to 50 mM NaCl. CPN60αβ1β2 containing fractions were concentrated and applied to Superdex200 gel filtration. Purified CPN60αβ1β2 eluted in the void volume. Purified CPN60αβ1β2 was concentrated to ~2 mg/mL, flash-frozen in liquid nitrogen, and stored at −80°C.

### Co-immunoprecipitation assays

For total protein extraction, *Chlamydomonas reinhardtii* CC-400 (*cw15 mt+*) was grown in 2 L TAP medium to a density of 5×10^6^~1×10^7^. Cells were harvested by centrifugation and resuspended in lysis buffer (20 mM HEPES-KOH pH 7.5, 150 mM NaCl, 10 mM MgCl_2_, 5 mM KCl, 1 mM EDTA and 1 mM PMSF). Cells were broken by sonication and loaded onto a sucrose cushion (lysis buffer with 1 M sucrose) and centrifuged for 30 min at 150000 g. Supernatant fractions were decanted and kept on ice for further use. For stroma protein extraction was dialyzed to the same lysis buffer as total protein before the co-immunoprecipitation experiment. Protein A-Sepharose beads with coupled Cpn20 antibodies were equilibrated in lysis buffer and incubated with total protein or stroma protein under agitation for 1 h at 4°C. Beads were washed three times with lysis buffer containing 0.1% Triton and once with 10 mM Tris-HCl pH 7.5. Bound protein complexes were eluted by incubation with 1×SDS loading buffer lacking β-mercaptoethanol by shaking for 1 h at 4°C. Eluted protein complexes were separated on a 5%-13% SDS-polyacrylamide gel. The key bands were cut out and proteins therein identified by LC-MS/MS.

### Pull-down assays

A total of 500 μg of CPN11_6xHis_2023 was immobilized on Ni-NTA beads and incubated with stroma proteins from 1 L cells for 2 h at 4°C in pull-down buffer (50 mM Tris–HCl pH 8.0, 10 mM MgCl_2_, 5 mM KCl, and 150 mM NaCl), supplemented with or without 1 mM ATP. Beads were washed three times with the pull-down buffer containing 40 mM imidazole. Proteins were eluted with pull-down buffer containing 250 mM imidazole, and 10 μl of the samples were resolved by SDS–PAGE and detected by immunoblotting.

### ATPase activity assays

ATPase activities of the chaperonins were measured spectrophotometrically using a coupled enzymatic assay. The assay was carried out in a mixture of 20 mM MOPS/KOH pH 7.5, 100 mM KCl, 10 mM MgCl_2_, 1 mM phosphoenolpyruvate, 20 units/ml pyruvate kinase, 30 units/ml lactate dehydrogenase, 0.25 mM β-nicotinamide adenine dinucleotide, reduced disodium salt hydrate (NADH). 0.5 μM co-chaperonin and 1 mM ATP were added, followed by incubating for 2 min at 25°C to remove any ADP present. The reaction was started by the addition of the 0.2 μM chaperonin complex, and the absorbance at 340 nm was monitored every 30 second over 10 min.

### Rubisco carboxylase activity assay

The Rubisco carboxylase activity assay was performed as described previously (Guo et al., 2015). Samples containing refolded Rubisco were mixed with 15 mM NaHCO_3_, 0.1 μCi /μl NaH^14^CO_3_, 20 mM MgCl_2_, and 0.2 mM DTT and incubated for 5 min. Next, 2.5 mM ribulose-1,5-bisphosphate (RuBP) was added into the sample mixtures to start carboxylation and incubated for 10 min. The reaction was terminated with 3 M acetic acid. The dried mixture was dissolved in 100 μl of ddH_2_O and mixed with 1 mL of scintillation fluid. The amount of fixed ^14^C was counted with a liquid scintillation counter (PerkinElmer, Waltham, MA, USA).

### MGM100 complementation assay

*E.coli* strain MGM100 was transformed with pQlinKT plasmids encoding the (co-)chaperonins by a standard electroporation procedure. A single colony was picked and inoculated in LB medium with 0.02% arabinose. Cells were collected by centrifugation when the culture reached an OD_600_ of 1.0, and washed five times using LB liquid medium with 0.5 mM glucose. The harvested cells were resuspended in 1 ml of LB and 10-fold gradient dilutions were made. Cell suspensions were then spotted onto an LB agar plates containing 0.5 mM glucose and 0.1 mM ITPG. Plates were incubated either at 37°C or 45°C for 12 h.

### RrRubisco refolding assay

Recombinant dimeric RrRubisco (Rubisco from *Rhodospirillum rubrum*) was denatured in denaturation buffer (6 M guanidinium hydrochloride, 10 mM DTT) for 1 h at 25°C, and then 1 μM denatured RrRubisco was diluted into ice-cold refolding assay buffer (20 mM MOPS-KOH, pH 7.5, 5 mM Mg(OAc)2,100 mM KCl, 5 mM DTT) containing 0.5 μM CPN60αβ1β2. The mixture was incubated for 5 min at 25°C for sufficient binding of denatured RrRbcL to CPN60αβ1β2. Unbound RrRbcL was removed by centrifugation for 10 min at 16,000 g. The supernatant was transferred into new tubes and supplied with 1 mg/ml BSA and 1 μM co-chaperonin complex. Refolding was initiated with 2 mM ATP and the reaction was stopped with 10 mM glucose and 2.5 U of hexokinase.

### Quantitative analysis of Cpn60αβ1β2 subunit stoichiometries

The isolated CPN60 holocomplex was separated by SDS-PAGE and by ND-PAGE. In addition, the QconCAT protein covering all Chlamydomonas CPN60 subunits reported previously by (Bai et al., 2015) was allowed to just migrate into an SDS-polyacrylamide gel. The proteins were stained with colloidal Coomassie G (Neuhoff et al., 1988) and bands containing CPN60 subunits and the QconCAT protein cut out followed by tryptic in-gel digestion as described previously (Muller et al., 2018). Three biological replicates were analyzed with two technical replicates by microliquid chromatography tandem mass spectrometry (μLC-MS/MS) using a TripleTOF 6600 instrument coupled to an Eksigent 425 HPLC system (AB SCIEX, Darmstadt) as described previously (Muller et al., 2018)with the following modifications: the HPLC gradient (buffer A: 2% acetonitrile, 0.1% formic acid; buffer B: 90% acetonitrile, 0.1% formic acid; flow rate of 4µl / min) ramped from 99 % buffer A, 1 % buffer B to 33% buffer B within 19 min, then to 50% buffer B within 3 min, followed by washing and equilibration steps. The mass spectrometer was operated in MRM-HR mode, i.e., one MS1 (m/z 350 – 1250, 300 ms accumulation time) was followed by MS2 scans of the selected precursor ions (m/z 100 – 1500, 30 ms accumulation time, collision energy as indicated in Supplementary file 1 resulting in a total cycle time of 1 s. Data analysis was performed using Skyline (v4.1.0.11796) (MacLean et al., 2010). The XICs of the four top ranked transitions per peptide were extracted (retention times, precursor m/z values and transitions are given in Supplementary file 1) and summed areas for the different proteins in the CPN60 samples were normalized to the respective areas of the QCONCAT protein (wherein a 1:1 ratio of all peptides is given). The resulting fractional abundances for the three CPN60 subunits were then converted to ratios assuming a 14mer complex.

### Cryo-EM sample preparation and data collection

To homogenize the CPN60αβ1β2 sample, purified CPN60αβ1β2 (about 2 mg/ml) was incubated in the presence of 10 mM MgCl_2_, 10 mM KCl and 2 mM ADP for 15 min at 25 °C before freezing. A 2 µl drop of samples was placed onto a glow-discharged Quantifoil holey carbon grid (R1.2×1.3, 200 mesh, Quantifoil Micro Tools). The grid was plunge-frozen in liquid nitrogen-cooled liquid ethane using an FEI Mark IV Vitrobot. Moreover, to prepare the CPN60αβ1β2-CPN11/20/23 complex sample, purified CPN60αβ1β2 and CPN11/20/23 were mixed at a molar ratio of 1:2 in the presence of 10 mM MgCl_2_, 10 mM KCl, and 2 mM ADP for 15 min at 30°C. The same sample vitrification process used for CPN60αβ1β2 was followed for CPN60αβ1β2-CPN11/20/23.

The frozen-hydrated sample was subsequently imaged in an FEI Titan Krios transmission electron microscope operated at 300 kV and equipped with a Cs corrector. Images were collected by using a Gatan K2 Summit direct electron detector in super-resolution mode with a pixel size of 0.65 Å, which were further processed after bined by 2 times, generating a final pixel size of 1.30 Å. Each movie was dose-fractioned into 38 frames. The exposure time was set to be 7.6 s with 0.2 s for each frame, producing a total dose of ~38 e^−^/Å^2^. Defocus values for this dataset varied from –0.8 to – 2.5 μm. All the images were collected by utilizing the SerialEM automated data collection software package(Mastronarde, 2005) .

### Image processing and 3D reconstruction

Image processing and reconstruction processes are provided in SFigure 6. For CPN6060αβ1β2, a total of 1,447 cryo-EM images were collected. All 38 frames for each movie were aligned and summed into a single micrograph using MotionCor2(Zheng et al., 2017) with bin by 2, the dose-weighted micrograph was used for further image processing. The CTF parameters were determined with CTFFIND4(Rohou and Grigorieff, 2015). Particles were picked automatically with RELION2.0 (Kimanius et al., 2016; Scheres, 2012), and bad particles and ice contamination were excluded by manual selection and 2D classification. Eventually, 140,954 particles were selected for further processing. We used the CPN60β1 crystal structure (PDB ID: 5CDI)(Zhang et al., 2016a) as the premier initial model for 3D classification, which was low-pass-filtered to 60 Å by using e2pdb2mrc.py program in EMAN2.1 (Tang et al., 2007). No symmetry was applied in our 3D classification process. One of the four classes with better structural features was selected as final initial model. Then another round of 3D classification was carried out. After that, we re-extracted and re-centered the cleaned-up 82,545 particles based on the refined coordinates, and did another round of 3D refinement and further post-processing in RELION2.0. We finally got a density map at the resolution of 4.06 Å based on the gold-standard Fourier shell correlation (FSC) 0.143 criterion(van Heel and Schatz, 2005).

For the CPN60αβ1β2-CPN11/20/23 complex, except when otherwise mentioned, the procedure used for CPN60αβ1β2 was adapted. A total of 2,808 cryo-EM images were collected. After manual selection and 2D classification, 189,854 particles were selected for subsequent processing. We used our CPN60αβ1β2 cryo-EM map but low-pass filtered to 50 Å as intial model. After 3D classification, one of the four classes apperaing with the CPN11/20/23 lid and with better structural features was selected as final initial model. Finally, the cleaned-up 67,808 particles yielded a density map at the resolution of 3.82 Å based on the gold-standard Fourier shell correlation (FSC) 0.143 criterion.

### Homology model building and analysis

To analyze the structural data in a more details, we built homology models of CPN60α, CPN60β1, and CPN60β2, respectively, by Modeller using GroEL subunit (PDB ID: 1AON) as the template (Marti-Renom et al., 2000; Webb and Sali, 2014). Since the *cis*-ring of CPN11/20/23-CPN60αβ1β2 was beter resolved than the *trans* ring or the CPN60αβ1β2 alone map, we than use this density to further refine the general conformaiton of the homology models. This *cis*-ring map appears highly symmetrical and the rotational correlation analoysis also revealed nearly identical conformations among the seven CPN60 subunits (Fig. 5A middle panel, Fig. 5B middle panel). We then placed the homology models of CPN60α, CPN60β1, and CPN60β2 as rigid-body into a randomly selected single-subunit density of CPN60 in this cis-ring using UCSF Chimera (Pettersen et al., 2004). The models were further flexiblely refined against the subunit density by Relax program in Rosetta software package and some regions were refined mannually in COOT (Emsley et al., 2010; Gray et al., 2003). The amino acid hydrophobicity and electrostatic potential were calculated in UCSF chimera and all structural images were generated by utilizing UCSF Chimera (Pettersen et al., 2004).

### Accession codes

Electron density maps have been deposited in the Electron Microscopy Data Bank under accession codes EMD-6947 for CPN60αβ1β2-ADP and EMD-6946 for CPN11/20/23-CPN60αβ1β2-ADP.

## Acknowledgments

We are also grateful to the staff of the NCPSS EM facility, Database and Computing facility for instrument support and technical assistance. This work was funded by the National Natural Science Foundation of China (31671262, 31670754), the National Basic Research Program of China (973 Program, 2015CB150106-1, 2017YFA0503503), and the Deutsche Forschungsgemeinschaft (TRR175).

## Author Contributions

C. L. and Y. C. supervised the design and interpretation of the project. Q. Z. executed most of the biochemical experiments. X. Z. collected the cryo-EM data and reconstructed the structure. Q. Z. and X. Z. performed the structural analysis. F. S. and M. S. performed the quantitative mass spectrometry. N. T. and N. W. are involved in the CPN60 complexes purification. Q. Z., X. Z., Y.C, and C. L. wrote the manuscript.

## Supplementary information

Supplementary data including 10 Figures and 1 file can be found enclosed with this article.

